# White matter differences in Parkinson’s disease mapped using tractometry

**DOI:** 10.1101/209858

**Authors:** Conor K. Corbin, Vikash Gupta, Julio E. Villalon-Reina, Talia M. Nir, Faisal M. Rashid, Sophia I. Thomopoulos, Neda Jahanshad, Paul M. Thompson

## Abstract

Neurodegenerative disorders are characterized by a progressive loss of brain function. Improved precision in mapping the altered brain pathways can provide a deep understanding of the trajectory of decline. We propose a tractometry workflow for conducting group statistical analyses of point-wise microstructural measures along white matter fasciculi to identify patterns of abnormalities associated with disease. We combined state-of-the-art tools including fiber registration, tract simplification and fiber matching for accurate point-wise statistical analyses across populations. We test the utility of this method by identifying group differences between Parkinson’s disease (PD) patients and healthy controls. We find statistically significant group differences in diffusion MRI derived measures along the anterior thalamic radiations (ATR), corticospinal tract (CST) and regions of the corpus callosum (CC). These pathways are essential for motor control systems within cortico-cortical and cortico-subcortical brain networks. Moreover, the reported pathological changes were not widespread but rather localized along several tracts. Point-wise tract analyses may therefore offer an advantage in anatomical specificity over traditional methods that assess mean microstructural measures across large regions of interest.

## 1. INTRODUCTION

Diffusion MRI (dMRI) is a non-invasive imaging modality that maps the diffusion of water molecules in the brain. dMRI has been used to detect alterations in white matter (WM) architecture in neurodegenerative disorders such as Parkinson’s [1] and Alzheimers disease [2]. Previously, white matter changes have been studied either by using tract based spatial statistics (TBSS) or by computing mean dMRI microstructural measures, such as fractional anisotropy (FA),mean diffusivity (MD), radial diffusivity (RD), axial diffusivity (AD) at different WM regions. In TBSS, a white matter skeleton is calculated and FA values along the skeleton are compared across groups of subjects. In another related work, diffusion metrics along the mean fiber tracts in different WM fiber bundles are compared [3]. Most of these methods discard information by analyzing diffusion metrics only along mean fiber tracts or the mean FA values of an ROI. Furthermore, voxelwise approaches are prone to errors either due to partial voluming or misregistration. [4]. In contrast to voxel based analysis, tractometry refers to a statistical procedure for comparing dMRI metrics along fiber tracts across populations. The method involves identifying corresponding fibers across subjects based on a predefined metric, projecting dMRI derived measures along fibers tracts followed by statistical analysis along the curves. There have been several tractometry studies in the past [3, 5, 6] that use such techniques, but significantly reduce the dimensionality of the data by analyzing mean diffusion values across an entire fiber bundle [5], or across a single mean fiber [3]. While these methods can be appealing due to their simplistic representation, they limit the ability to study variations across the full set of fibers in a bundle. White matter fiber bundles are large, dense structures that exhibit a wide range of microstructural properties [7]. To study the variability of these measures along the entire bundle, a template corresponding to each bundle is built. This template is designed to be a statistical representation of the the population. Based on a distance metric, fiber tracts corresponding to the ones in the template are found in all individuals and compared. Here, we describe a tool to perform all necessary steps for tractometry, and provide a real-case example to capture differences between patients with Parkinsons disease (PD) and age-matched controls point-wise along corresponding fibers in bundles representing white matter fasciculi.

PD is a neurodegenerative disorder characterized by motor function loss, muscle rigidity, and tremors [1]. These effects are partially caused by a loss of neurons in substantia nigra. This loss inhibits the transmission of dopamine, which is used to balance signal flow in motor control pathways. In PD, we expect to observe microstructural changes along fiber bundles connecting the motor cortex to other cortical and sub-cortical regions, including the thalamus. We use the proposed tractometry workflow to automatically extract bilaterally in all subjects the anterior thalamic radiations (ATR), superior longitudinal fasciculi (SLF), corticospinal tracts (CST), six regions of the corpus callosum (CC) and the fornix. We conduct statistical analysis to test the differences in WM microstructure between patients and controls throughout each bundle. As mentioned earlier, there have been several methods in the past that allow comparisons of diffusion metrics along white matter bundles. The methods presented in this paper compare diffusion measures along fiber bundles in the subject space, thus mitigating the effects of voxel-based registration errors as in the case of TBSS based methods.

## 2. METHODS

### 2.1 Data and Preprocessing

Analyses were performed on the publicly available Park-TDI dataset [1], (https://www.nitrc.org/projects/parktdi/). Park-TDI contains dMRI scans from 27 PD patients (ages: 50-81; 14M/13F) and 26 age matched controls (ages:47-78;14M/12F). The dMRI scans consist of 120 directions in each of two diffusion weighted shells b=1000, 2500 mm/sec2. Generalized Q-sampling imaging reconstruction was used for whole brain deterministic tractography using DSI-Studio with a max fiber count set to 400,000 (http://dsistudio.labsolver.org) A generalized fractional anisotropy (GFA) map was computed from the Q-sampling reconstruction, and tensor derived measures, i.e fractional anisotropy (FA), mean diffusivity (MD), radial diffusivity (RD), and axial diffusivity (AD), were computed with DIPY after fitting a weighted least squares tensor reconstruction (http://nipy.org/dipy/).

### 2.2 Fiber Clustering

For each subject, WM tracts were automatically extracted by querying fiber tracts that intersected particular regions of interest (ROIs), as shown in Figure 1b. ROI definitions were derived from the Type II JHU-EVE Atlas [8]. The EVE atlas was registered to each subjects FA map using the ANTS nonlinear registration [9]. The resulting deformation field was used to warp the ROI labels to each subjects FA image using nearest neighbor interpolation. We extracted the left and right SLF, left and right (ATR), the full corticospinal tract (CST) and fornix. Further we extracted the corpus callosum (CC), and partitioned it into fiber bundles connecting the frontal, temporal, parietal, and occipital regions, as well as regions connecting pre and postcentral gyri.

**Fig. 1.**
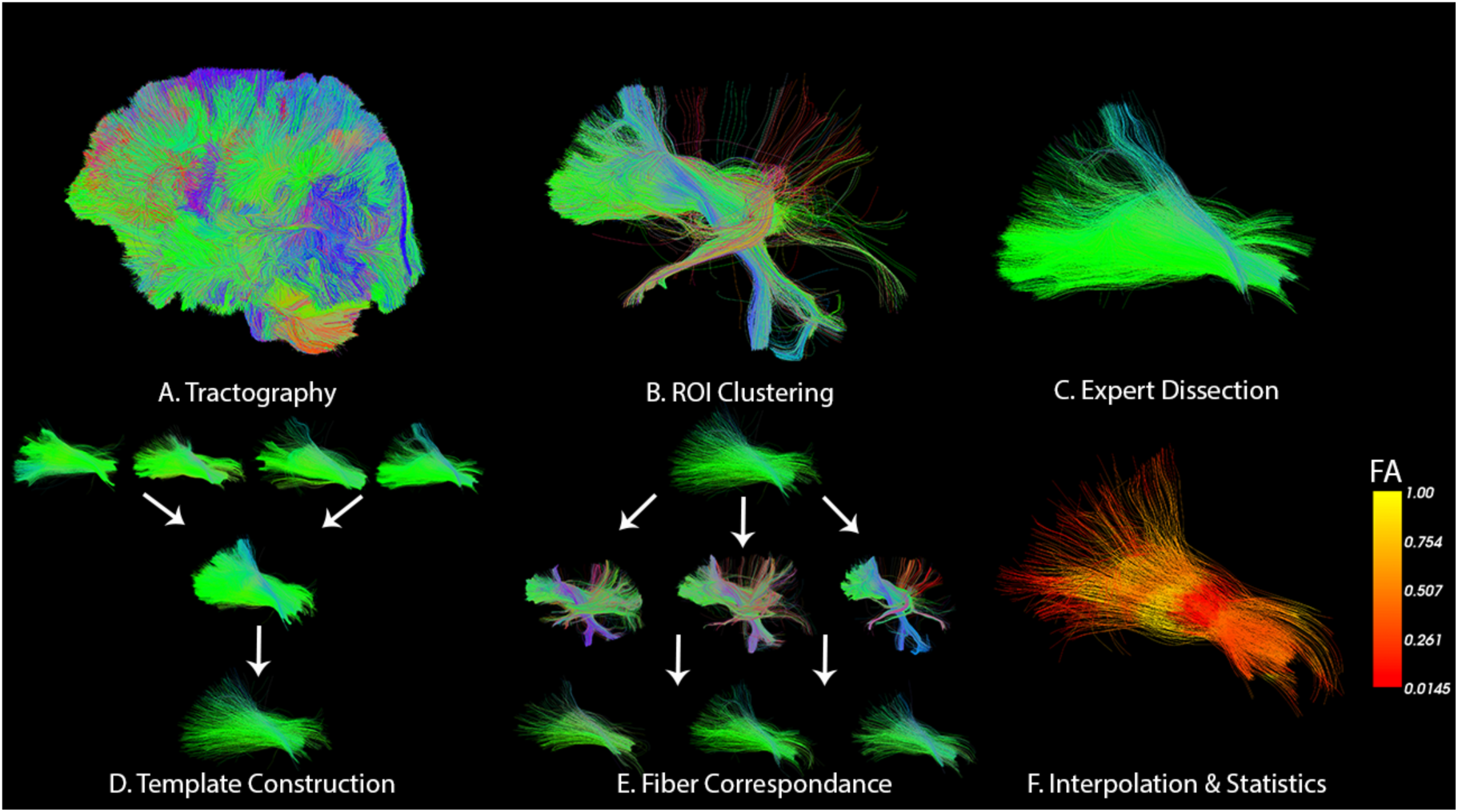
Major steps involved in our tractometry pipeline. A) Tractography is computed from preprocessed DWIs. B) Tractograms are segmented into bundles with ROI based clustering. C) Four subjects are randomly chosen, and bundles are cleaned to remove spurious fibers. D) Cleaned bundles are merged and centroids are extracted to create a template. E) The template is used to find corresponding fiber tracts across all of the subjects. F) Diffusion metrics are interpolated onto the corresponding fiber tracts and group analysis is performed.

### 2.3 Template Construction

After the ROI based fiber clustering, there remained a considerable number of false-positive fibers tracts in the extracted fiber bundles. False-positives in four randomly selected subjects were removed by a neuroanatomical expert. These “cleaned” fiber bundles from the four subjects serve as the basis for population templates bundles used in subsequent analyses. Anatomically corresponding bundles across the four subjects were rigidly registered using DIPY’s Direct Streamline Registration (DSR) tool [10], and merged to one set of fiber tracts. In each of these merged sets, a representative subset was extracted (centroids) using DIPY’s Quickbundles (QB) algorithm [11]. We define this subset of fibers for each fasciculi as population template bundles (P-bundles). We performed a gradient descent on the QB distance threshold to optimize the number of centroids produced so that each P-bundle contained a similar number of fiber tracts. The template construction is shown for the left anterior thalamic radiation in Figure 1D.

### 2.4. Fiber Correspondence

All fibers tracts were sampled to contain an equal number of points. For computing the correspondence between fibers tracts across subjects, we utilized the minimum average direct flip (MDF) distance metric, which is also used in the QB algorithm [11]. Using this metric, we matched each fiber tract in the P-bundles to a fiber tract in the subjects corresponding tract bundle (extracted with ROIs) by choosing the closest fiber. If the closest fiber tract resided outside of a 10mm distance, the correspondence was labeled null. Often, multiple P-bundle fibers pass through a single voxel. Thus, for a given voxel with multiple fiber points, we calculated a mean of the interpolated scalar measures to avoid multiple comparisons. All fiber tracts are oriented such that the first point in a subject fiber corresponds to the first point in the corresponding P-bundle fiber tract. Fiber correspondence is visually represented in Fig 1E.

### 2.5. Statistical Analysis

Multiple linear regression analysis was conducted to test for case/control group differences in microstructural measures point-wise along each fiber bundle. We controlled for age, sex and intracranial volume (ICV) and corrected for multiple comparisons across each fiber bundle using the false discovery rate (FDR) procedure at a threshold of q=0.05.

## 3. RESULTS

Compared to healthy controls, PD patients showed significant localized differences in WM microstructure in ten of the thirteen bundles (Table 2), with findings largely localized to callosal interhemispheric connections of the postcentral gyrus and the parietal and occipital lobes. Pointwise FDR-corrected p-values across streamlines within template bundles are shown in Figure 2. PD patients showed lower FA and GFA, and increased MD, RD in these regions. We also detected higher FA and GFA in the anterior thalamic radiations bilaterally. Increases in FA in the PD population compared to controls have been reported before [3].

**Fig. 2.**
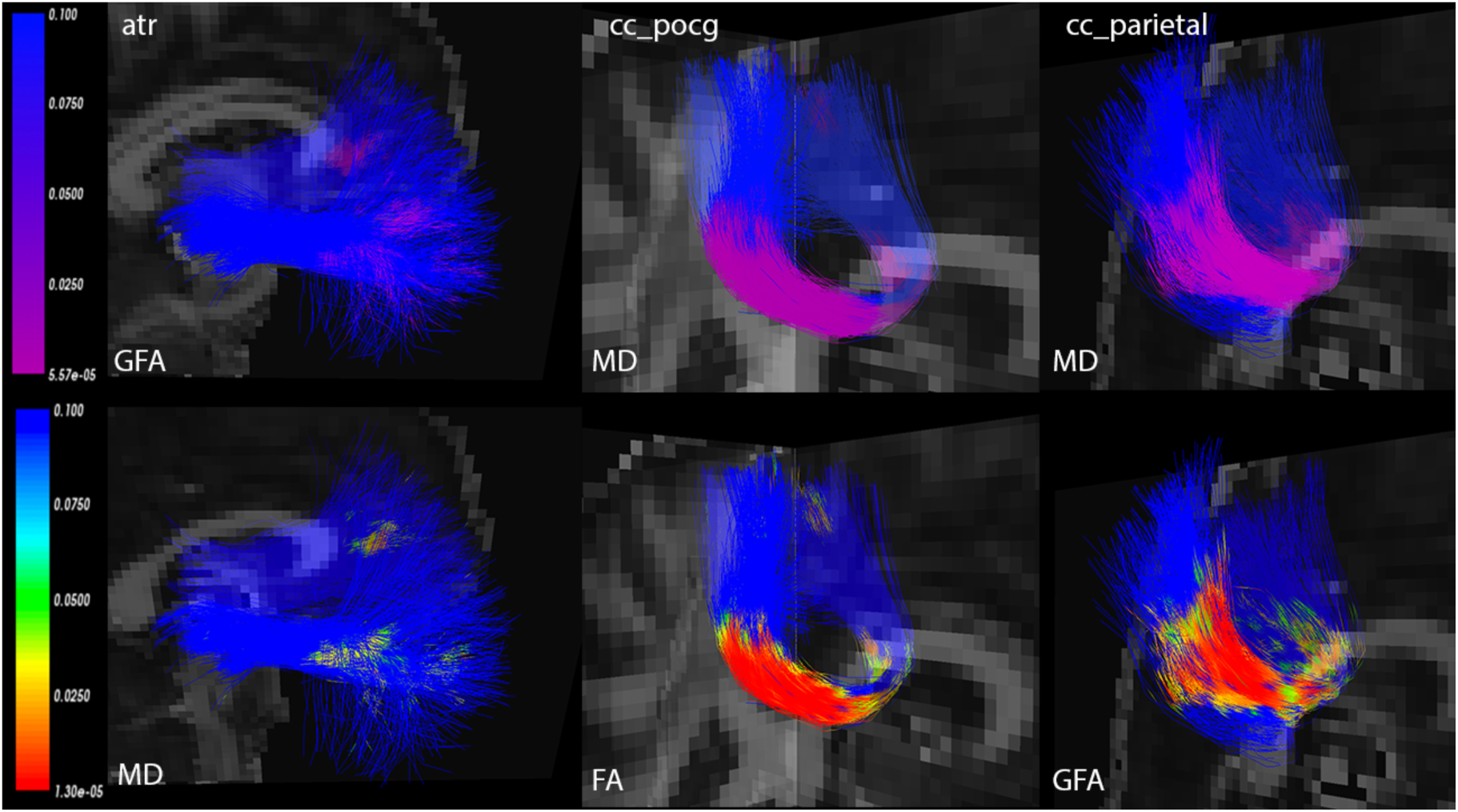
Corrected p-values projected onto the P-bundles showing regions with statistically significant differences between PD patients and controls. Purple signifies greater metric values in PD compared to controls, while green to red signifies lesser metric values. P-values were corrected with the false discovery rate threshold at an alpha of 0.05.

**Table 1.**
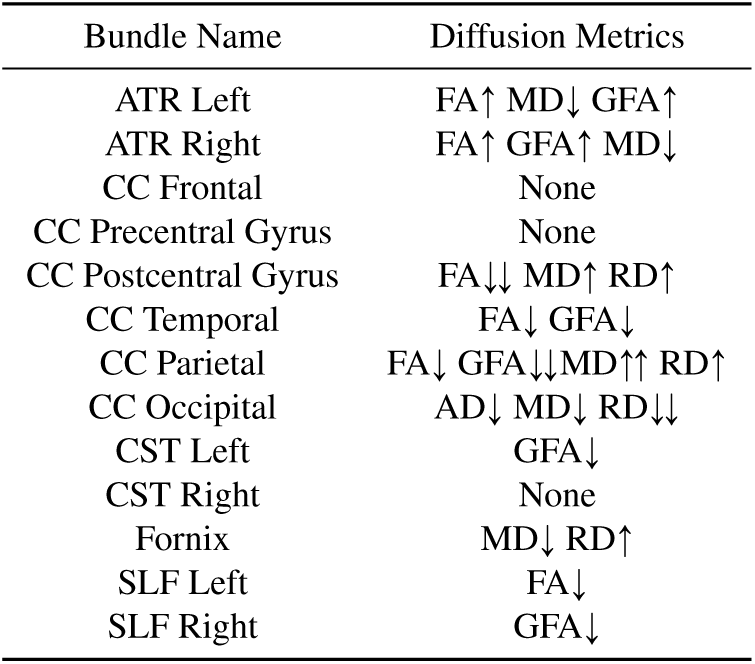
Bundles with statistically significant differences in the following diffusion metrics. The arrows represent the direction of the difference compared to the healthy population. Double arrows signify that at least 15 percent of the tests in the bundle passed the multiple comparison test.

## 4. CONCLUSIONS

Here we present an entire workflow for conducting statistical analysis of WM microstructure along streamlines within major fasciculi. The analysis, highlighted here for its promise for Parkinsons disease research, allows users to assess potential deficits along the extent of a given WM tracts, as opposed to summarizing the differences into a single measure. In summary, our workflow builds off of widely tested and validated open-source tools to develop a novel population based analysis technique, combining anatomically driven bundling of tractography derived streamlines, subject-wise correspondence, and mapping of dMRI microstructural properties along the corresponding points for population based statistics, complete with multiple comparisons correction. The methods presented in this paper can be applicable to study any disease or characteristic pertaining to white matter.

One of the limitations of the proposed workflow is the relatively low number of subjects used for creating the template bundles. Creating fiber bundle templates requires manual segmentation because of the huge number of spurious fiber tracts present in the tractogram, thus making the process expensive. However, in the future, we intend to use FiberNet [12], a deep learning tool for automatic extraction of fiber bundles. Using this tool, it will become feasible to construct a population template using a larger sample of individuals.

